# Informative priors contribute to quantifying the occurrence rate of a rare tree-related microhabitat in a managed forest

**DOI:** 10.1101/2024.11.28.625900

**Authors:** Pierre Cottais, Benoît Courbaud, Laurent Larrieu, Nicolas Gouix, Fabien Laroche

## Abstract

Tree-related microhabitats (TreMs) are key features for forest biodiversity, and knowing their accumulation rate is essential to design integrative management strategies. Many types of TreMs are associated to large old trees and show slow ontogenical processes. The rarity of such TreMs (particularly in intensively managed forests) hinder the estimation of their occurrence rate along tree growth. Here, we used a continental meta-analysis on TreMs occurrence rate along tree growth to build informative priors for a model of trunk-base rot-hole occurrence on oaks within the Grésigne forest, France — a context where stand management and tree DBH were confounded. We explored whether the use of informative priors could improve the identifiability, the precision of estimates and the predictive abilities of the model. Without prior information, the low variance of tree DBH within management modalities rendered the model poorly identifiable and prevented the detection of an effect of tree DBH *per se* across the range of explored tree DBH. By contrast, using informative priors contributed to improve the precision of estimates and lead to detecting a positive effect of tree DBH *per se*. Informative priors did not degrade the model fit and clearly improved predictive abilities on new stands. In particular, while the model without prior information did not predict the occurrence of trunk-base rot-holes significantly better than a purely random guess, the model with informative priors did. Irrespective of the prior used, models suggested that the high recruitment of trunk-base rot-holes in Grésigne may be a temporary management effect in stands undergoing conversion from coppice-with-standards to high forest through sprout thinning, which will lead to conservation issues for cavicolous saproxylic species when all conversions are complete. Because using informative priors was simple and beneficial in our study, it should be further explored in other local applied contexts to orientate forest management.

## 1 Introduction

Some ecological processes are hard to estimate from individual empirical studies. It happens, for instance, when they are associated to rare events, like quantifying the rate of microhabitats formation on trees in the field of forest ecology. Tree-related microhabitats (TreMs) are tree morphological structures that constitute particular and essential substrates for thousands of species [Larrieu et al., 2018]. TreMs thus contribute to forest biodiversity, with heterogeneous effects across forest taxa and across geographical regions [Bouget et al., 2014, Paillet et al., 2018]. Some types of TreMs are inherently rare, because they are caused by rare natural events (e.g lightning scars on trunks). Other types of TreMs have slow ontogenical processes, which make them naturally associated to large, old trees. Short sylvicultural cycles make these types of TreMs rarer and more ephemeral in intensively managed forests than would be expected if unharvested [Vuidot et al., 2011]. The human-induced rarity of TreMs associated to old trees raises conservation issues for associated species, and calls for a better integration of TreMs in a multifunctional approach of forest management e.g. through keeping retention trees in harvested stands [Gustafsson et al., 2012] or extending the rotation length of forest stands [Percel et al., 2018]. Calibrating this type of measures requires quantifying the rate at which target TreMs occur on trees. Unfortunately, TreMs rarity hinders the quantification of their rate of occurrence within a target management site.

Mutualizing the information from many empirical studies is one way out of the difficulty of making inferences from rare events. Meta-analyses follow this rationale [Hobbs and Hilborn, 2006], combining multiple experimental results to synthesize knowledge about some target parameters and their variation among studies. The study of the rate of occurrence of TreMs across forest sites at continental scale by Courbaud et al. [2022] could be classified in this cate-gory. The authors derived global estimates of occurrence rates for several TreM types, and their analysis revealed a high spatial variance of these rates across sites included in the analysis (see their appendix S6a), in both managed and unmanaged forests. The spatial variance of TreMs occurrence rate could partially stem from the fact that TreM occurrence rate was quantified as a function of tree diameter but overlooked tree age, which is harder to obtain on a large scale basis. Tree diameter being kept constant, many TreM types are known to occur more often on older trees [Kozák et al., 2023]. Therefore any variation in tree growth speed (e.g. due to fertility) can create distortions across sites on the relationship between TreMs occurrence and diameter. In a meta-analysis across France, Germany, Slovakia and Romania, Spînu et al. [2024] could avoid this ambiguity by considering explicit temporal data, however the time range of their study lead them to focus on the phenomenon of losing ephemeral TreMs in time rather than recruiting TreMs with slow ontogeny. The spatial variance of TreMs occurrence rate can also stem from differences in management that can modulate the rate of occurrence of some TreMs, e.g. those induced by damages during sylvicultural interventions such as bark loss along skidding tracks. More generally, specific processes leading to a TreM can vary a lot among sites (e.g. rockfalls and strong slopes specifically contribute to TreMs creation in moun-taneous sites). Overall, the presumably strong variation among forest sites of TreMs occurrence rates questions how global studies like Courbaud et al. [2022] or Spînu et al. [2024] can be used by managers to improve inferences of a TreM occurrence rate within one target forest site.

Bayesian models offer the opportunity to use the information contained in previous studies within a new experiment involving the same processes of interest, under the form of ‘informative priors’ on associated parameters [McCarthy and Masters, 2005, Clark, 2007]. Informative prior distributions can be ob-tained from similar empirical studies [Ellison, 2004, Dupuis and Joachim, 2006] or from data generated in pilot experiments [Morris et al., 2013]. They also allow transferring information about target parameters across studies that use different types of data, designs etc. as long as they share parameters that points to the same processes. This has been used to learn from observational data in controlled experiments [McCarthy and Masters, 2005], to bridge gaps in time series of long term monitoring programs [Rodhouse et al., 2019], to generate prior for bird mortality rate based on their bodymass [McCarthy and Masters, 2005], to generate prior for tree mortality based on their growth speed [Morris et al., 2015]. However this has little been applied to the use of large-scale information in specific local case-studies, especially not for the quantification of TreM occurrence dynamics in forest.

Like any regularization procedure [Hooten and Hobbs, 2015], the use of informative priors in Bayesian analyses mechanically improves the precision of estimates (i.e. they decrease the variance of posterior distribution of parameters; McCarthy and Masters [2005], Garrard et al. [2012], Morris et al. [2013]). However, because it comes from other studies, the prior information may be partially unadapted to the current context, and the increase in precision of estimates may come at the cost of larger estimation bias leading to lower predictive accuracy. In their study on tree mortality with informative priors based on tree growth rate, Morris et al. [2015] showed that while the precision of estimates was systematically improved compared to non-informative priors, predictions were improved for only half of the species. Beyond the need of building informed priors based on a repeatable and transparent procedure, it is thus necessary to use goodness of fit metrics and validation procedures to assess the potential loss of accuracy [Hooten and Hobbs, 2015, Gelman and Hennig, 2017, Banner et al., 2020].

Here, we aimed at using the information of the continental analysis by Courbaud et al. [2022] on TreMs occurrence rates across Europe and Iran to study the dynamics of one specific TreM type within a single forest, using Bayesian informed priors. We focused on the specific case of trunk-base rot-holes (Fig. 1) in the Grésigne forest, south-west France. In this case study, oldest stands were undergoing conversion from coppice-with-standards to high forest by singling, a treatment known to positively affect the prevalence of trunk-base rot-holes [Gouix, 2011]. By contrast youngest plots were managed as regular high forest. As a result, stands managed as regular high forest tended to harbour oak trees with a lower and narrower range of diameters than stands managed as high forest by singling. One thus expected a confounding effect between tree diameter and stand management effects on the occurrence of trunk-base rot holes. We aimed at illustrating how the use of informative prior could improve the identifiability and the precision when estimating the effects driving the rate of occurrence of trunk-base rot-holes in Grésigne. We also aimed at quantifying the potential biases and loss of predictive accuracy induced by the use of informative priors. We thus hoped to clarify whether this technically simple procedure can generate better local information on TreMs, hence opening the way to designing well-adjusted multifunctional forest management strategies.

**Figure 1:**
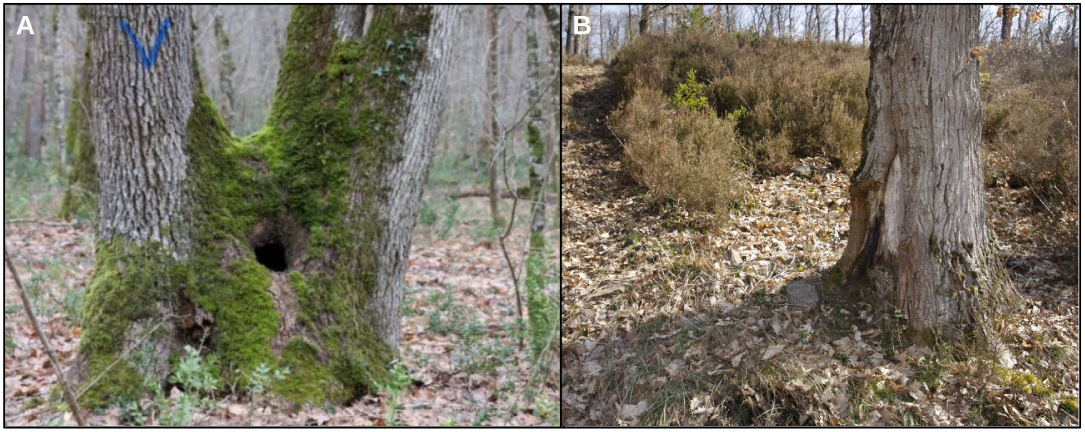
Examples of trunk-base rot-holes on oak trees within Grésigne forest. The trunk-base rot-hole in panel A likely comes from sprout thinning. The trunk-base rot-hole in panel B is located along a skidding track and likely comes from logging damages.

## 2 Material and methods

### 2.1 Oaks and trunk-base rot-holes data in Grésigne

We focused on one TreM type, trunk-base rot-holes, which corresponds to the category ‘trunk base rot-hole’ in the typology of Larrieu et al. [2018]. Trunk-base rot-holes harbour a chamber that is closed on top and completely protected from surrounding microclimate and rain. They contain wood mould, the quantity of which increases with the development stage of the hole. The bottom of the hole is in contact with the ground although the hole entrance can be higher on the trunk.

We surveyed trunk-base rot-holes in the Grésigne forest, a State forest of south-west France, in the Tarn department. It is a lowland forest, covering c.a. 3500 ha, and dominated by sessile oak (*Quercus petraea*). The forest has undergone a gradual conversion of management from coppice-with-standards to high forest, initiated in late XIX^th^ century. As a result, stands are currently managed in a heterogenous way. Oldest stands (more than 120 years since the last final cut) are still undergoing conversion from coppice-with-standards to high forest. In these stands, the last final cut corresponded to the singling operation that initiated the conversion process. In youngest stands (less than 120 years since the last final cut), the conversion has ended and a new silvicultural cycle has been initiated. These stands are thus managed as classic high forest and the last final cut corresponded to a seed tree removal cut. The conversion from coppice-with-standards to high forest by singling has contributed to generate a high occurrence of rot-holes at the bottom of oak trunks within stands undergoing conversion, particularly large holes at advanced development stage (*sensu* Larrieu et al. [2018]). As a result, the Grésigne forest harbours an outstanding biodiversity associated to such trunk-base rot-holes, including the violet click beetle *Limoniscus violaceus* which is listed in the second annex of the Habitat Directive [Gouix et al., 2015].

Nine 1-ha circular plots (i.e. radius of 57 m) were surveyed in 2021, six being located within stands that are still under conversion, and three within stands where conversion was complete and a new sylvicultural cycle of high forest has been initiated (Figure 2). These plots were part of the BloBiForM project, which aimed at generating a fractal sampling design [Laroche, 2022] to study *β*-diversity among ecological communities inhabiting trunk-base rot-hole over a range of spatial scales from tree to forest. All the trees within plots were inspected by two operators. For trees with diameter at breast height (DBH) above 17.5 cm, operators noted the tree species, the tree DBH and whether the tree harboured a trunk-base rot-hole. The data obtained during this first campaign was used to calibrate the local model of trunk-base rot-hole occurrence below.

**Figure 2:**
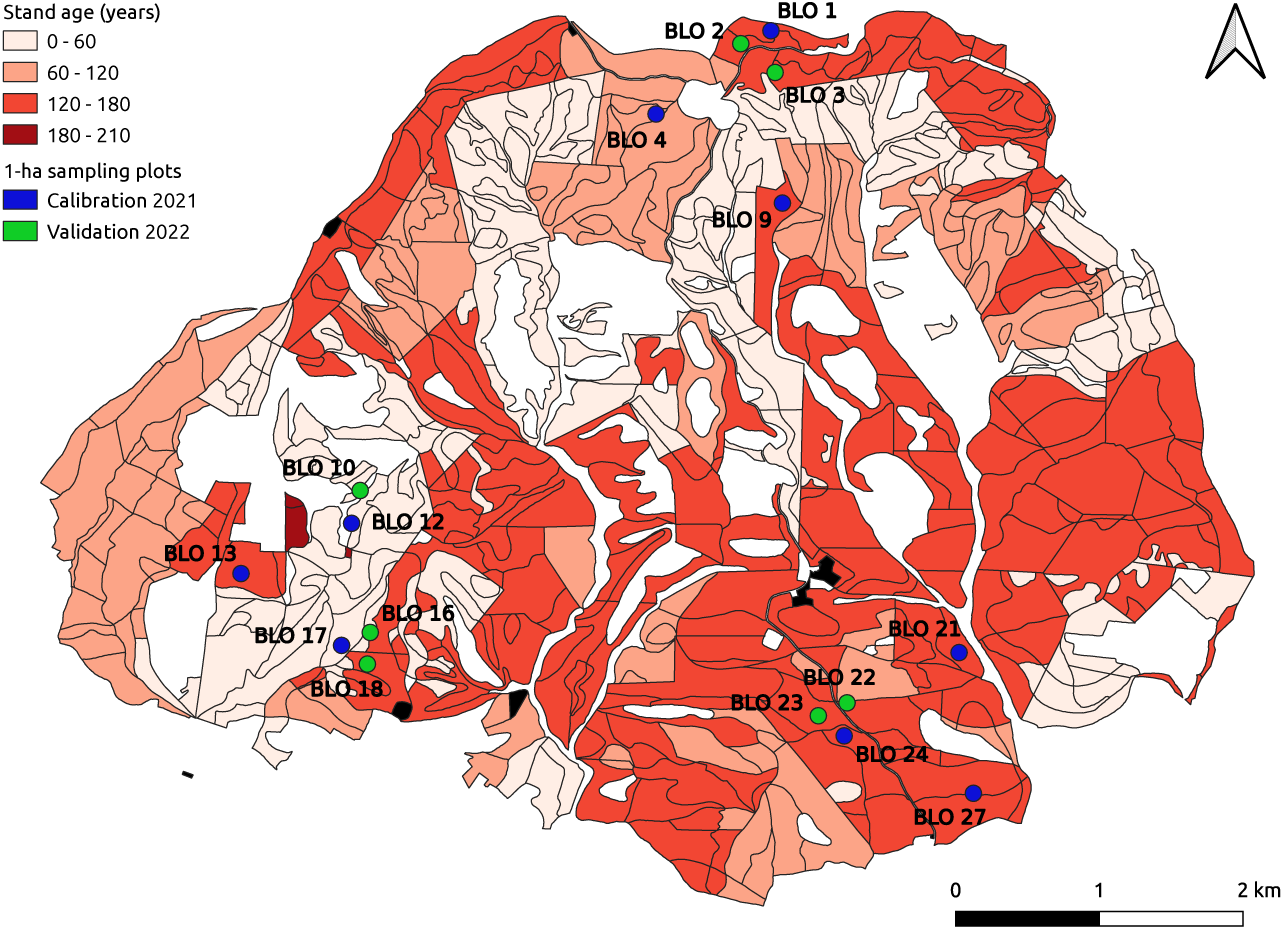
Study sampling design within the Grésigne forest. Circles present the sampling plots of 2021 (blue) and 2022 (green) surveys. Red shapes show the oak stands of the forest, and darker red indicates older stands (see time scale on the left). White areas indicate either non-forest areas (e.g. roads or forest outside) or non-oak forest stands. Black shapes correspond to buildings (e.g. the forest guards office).

Seven additional plots were surveyed in 2022 (Figure 2). Four plots were located in stands under conversion, one in a stand where conversion was complete. The two remaining plots were overlapping the two types of stands, but one was predominantly located in a stand undergoing conversion (*BLO*16) while the other was predominantly in a stand where conversion was complete (*BLO*22). Tree sampling occurred according to the same protocol (circular plots, two operators, DBH above 17.5 cm etc.). This new data was used as a validation dataset below. Although a fraction of the Grésigne forest (c.a. 40 ha) is a protected left-aside area, the 9 + 7 = 16 sampled plots of our study were all located in managed stands.

### 2.2 Continental model of trunk-base rot-hole occurrence

Courbaud et al. [2017] suggested to model the occurrence of TreMs on trees in forest as a Bernoulli random variable, using the framework of survival analysis. This framework relies on defining a survival function *S*(*d*) corresponding to the probability that a tree has not developped a TreM of a given type yet when reaching a DBH *d* (expressed in cm). They used a Weibull distribution to model survival functions:

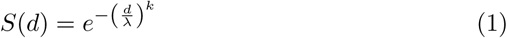

where *λ >* 0 is a scaling parameter of the survival function in terms of tree DBH. More precisely, *λ* is the tree DBH at which the probability of having developped a TreM of target type becomes higher than 1 *−* exp(*−*1) *≈* 63%. Parameter *k >* 0 drives the shape of the survival function. If *k >* 1, the rate of TreM formation on non-bearing trees increases with the DBH. Conversely, if *k <* 1, the rate decreases with the DBH. Courbaud et al. [2022] conducted a Bayesian estimation of the survival function parameters for 11 types of TreMs based on the observation of 80,099 living trees from 19 species groups in plots spread across 307 forests of Europe and Iran [Courbaud et al., 2021]. The authors built a separate model for each TreM type. The scale parameter of the survival function depended on tree species, coarse management regime (i.e. ‘managed’ versus ‘unmanaged’) and forest site. Precisely, for a given tree species, the scale parameter for a target TreM in forest site *s* with management regime *m* was expressed as:

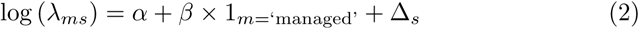

The term 1*_m_*_=‘managed’_ equates 1 when the tree is in a managed stand, 0 otherwise. The forest site effect Δ*_s_* was modeled as a random effect following a normal distribution with mean 0 and variance *σ*^2^. The shape parameter *k* of the survival function for oak trees was estimated through the log parameter *θ* = log(*k*). Parameters *α* and *θ* were specific to tree species, while *β* and *σ*^2^ were common to all tree species.

The TreM typology used by Courbaud et al. [2022] lumped all types of rot holes containing wood mould together in a single category including ‘Trunk base rot hole’ (the type of interest here), but also ‘Trunk rot hole’, ‘Semi-open trunk rot hole’, ‘Chimney trunk base rot hole’, ‘Chimney trunk rot hole’ (see Larrieu et al. [2018] for a description of these items). We therefore re-used their methods and code and model to generate sampled posterior distribution of parameters *α*, *β*, *θ* and *σ*^2^ for trunk-base rot-holes only. We only considered the ‘Quercus’ species group, which reduced the dataset to 8684 trees in 146 forests. Courbaud et al. [2022] approach involved using a MCMC approach through a Gibbs sampling algorithm provided by the package *rjags* [Denwood, 2016]. For each estimation, four Markov chains with independent initialization were implemented. A 5 *×* 10^3^ steps burning period was applied, followed by 7 *×* 10^3^ steps where parameter values were recorded every 15 steps, resulting in 500 samples in each chain. The quality of the convergence of the Markov Chains was assessed by checking whether the Potential Scale Reduction Factor index (PSRF) was less than 1.1 for all parameters and that the Monte-Carlo standard errors were less than 5% of the posterior mean of the parameters [Denwood, 2016]. In what follows, we approximated the posterior distribution of the four continental parameters *α*, *β*, *θ* and *σ* by a multivariate normal distribution with mean and variance-covariance matrix equal to the corresponding empirical estimates derived from the 4 *×* 500 = 2000 posterior distribution samples.

### 2.3 Local model of trunk-base rot-hole occurrence

The process of trunk-base rot-hole occurrence on oaks in Grésigne was also modelled through a Weibull survival function based on tree DBH. We modelled the scale parameter *λ_ip_* of the survival function for tree *i* in sampled plot *p* as:

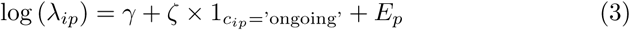

where we defined a ‘conversion’ covariate *c_ip_*, equal to 1 if the conversion towards high forest was still in progress for the stand where tree *i* of plot *p* lied (and current treatment is thus high forest by singling), and to 0 if the conversion was over. Parameter *ζ* thus corresponded to the effect of being in a stand under ongoing conversion. In general, all the trees within a plot belonged to the same stand, and thus had the same value of *c_ip_*. The plot effect *E_p_* was modelled as a random effect following a normal distribution with mean 0 and variance *τ* ^2^. The shape parameter *l* of the survival function was constant for all trees. When performing estimation, we considered the parameter *κ* = log(*l*) rather than *l* itself.

### Potential identifiability issues

— In Grésigne forest, oak trees are even-aged within stands, and stands where conversion has ended are younger than stands where conversion is still ongoing. From a statistical perspective, we thus expected the variance of tree DBH to be low within each management modality (i.e. conversion is ongoing or conversion has ended) and high between the two modalities. In this setting, the estimation of management and DBH effects becomes challenging. In particular, as explained in online supporting information [Laroche, 2025], we expect the region of parameters space defined by the following equations to be weakly identifiable :

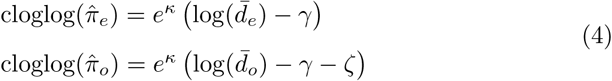

where where cloglog stands for the complementary log-log function cloglog(*p*) = log(*−* log(1 *− p*)), 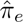 (resp. 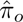) is the empirical proportion of trees with a trunk-base rot-hole within stands where conversion has ended (resp. where conversion is ongoing), and *d̅_e_* (resp. *d̅_o_*) is the average tree DBH within stands where conversion has ended (resp. where conversion is ongoing).

### 2.4 Bayesian estimation of local model parameters

Using a Bayesian framework for the estimation required to define priors for the four parameters *γ*, *ζ*, *κ* and *τ*^2^. We considered two alternative strategies, one with non-informative priors, the other with informative priors based on the posterior multivariate normal distribution of the continental model parameters. In both strategies, the prior distribution of the scale parameter of trunk-base rot-holes survival function was assumed identical between stands undergoing conversion and stands where conversion was over. Technically, this meant that prior distributions of *γ* and *γ* +*ζ* were set identical and independent by adjusting the prior distribution of parameter *ζ* [Laroche, 2025]. The prior on *τ* was set as a vague, uniform distribution between 0 and 20, independent from other parameters prior distribution. The two strategies differed in the aspects detailed below.

### Non-informative priors

— We used a vague prior with normal distribution, mean 0 and variance 10^2^ for parameter *γ*. Identical prior distribution for *γ* + *ζ* was obtained by setting a normal prior distribution for *ζ* with mean 0 and variance 3 *×* 10^2^, while setting the covariance between *ζ* and *γ* at *−*1.5 *×* 10^2^ [Laroche, 2025]. We used a vague prior with normal distribution, mean 0 and variance 10^2^ for parameter *θ*, and we assumed its independence with respect to *γ* and *ζ* priors.

### Informative priors

— We defined the prior distribution for *γ* as the posterior distribution of *α* + *β* +Δ in the continental model, where Δ was an independent normal distribution with mean 0 and variance *σ*^2^. In practice, the distribution was computed as a normal distribution with mean *m*(*γ*) = *M* (*α*) + *M* (*β*) and variance *v*(*γ*) = *V* (*α*) + *V* (*β*) + *C*(*α, β*) + *M* (*σ*^2^), where *M* (.), *V* (.) and *C*(., .) respectively stand for mean, variance and covariance of parameters in the continental model posterior distribution. Identical prior distribution for *γ* + *ζ* was obtained by setting a normal prior distribution for *ζ*, with mean 0 and variance 3 *× v*(*γ*) while setting the covariance between *ζ* and *γ* at *−*1.5 *× v*(*γ*) [Laroche, 2025]. We directly used the posterior normal distribution of *θ* derived from the continental model as a prior for *κ* in the local model.

We derived a sampled posterior distribution of parameters *γ*, *ζ*, *κ* and *τ*, in either case using a MCMC approach. We used a Gibbs sampling algorithm provided by the package *rjags* [Denwood, 2016], on the *R* software platform (v4.3.3; R Core Team [2023]). For each estimation, we implemented four Markov chains with independent initialization. We applied a 10^5^ steps burning period, followed by 2 *×* 10^6^ steps where parameter values were recorded every 2.5 *×* 10^3^ steps, resulting in 800 samples in each chain. The quality of the convergence of the Markov Chains was assessed by checking whether the Potential Scale Reduction Factor index (PSRF) was less than 1.1 for all parameters and that the Monte-Carlo standard errors were less than 5% of the posterior mean of the parameters [Denwood, 2016]. Codes and diagnostics are provided in a public repository online [Cottais and Laroche, 2024].

### 2.5 Goodness-of-fit of models

We first assessed the goodness of fit of each calibrated model through posterior predictive checks. For each parameter value in the sampled posterior distribution, we performed a simulation of the presence-absence of trunk-base rot-holes on all trees of the calibration dataset, hence generating 3200 virtual datasets for each model. We recorded three types of statistics: the number of trunk-base rot-holes in each stand, the total number of trunk-base rot-holes on oaks with DBH below 35 cm (median DBH of the dataset) and the total number of cavities on oaks with DBH above 35 cm. We assessed whether the values of statistics in the real dataset was comprised within a likely range of simulated values. To do so, we computed a bilateral p-value based on the empirical quantile of the observed value within simulated values.

We compared the goodness of fit of calibrated models using the Deviance Information Criterion (DIC; Spiegelhalter et al. [2002], McCarthy and Masters [2005]), which accounts for the fact that using informative priors decreases the effective number of degrees of freedom used by the model. Deviance is defined as *−*2 log(ℒ) where ℒ is the likelihood of the model upon the data. The DIC is computed as 2 *× D̅* (**Θ**) *− D*(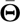), where *D̅* (**Θ**) is the mean deviance computed across the posterior distribution of parameters **Θ** and *D*(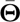) is the deviance at the mean posterior estimates of parameters. Means are computed empirically from the sampled posterior distributions. To compute DIC indices, we used **Θ** = (log(*γ*), *κ, ζ, E*_1_*, …, E*_9_). A lower DIC by more than two points is classically considered as a signal of significantly better fit.

### 2.6 Predictive performance of models

We used the 2022 dataset to assess the predictive performance of the two models calibrated in previous sections. For each model, we built upon the 3200 values of the sampled posterior distribution of parameters obtained from calibration on the 2021 dataset. For each parameter sample, we computed the predicted probabilities of presence of trunk-base rot-holes in all the trees of the validation dataset. The plot effect (*E_p_* in eq. (3)) was set to 0. In the two plots overlapping stands with distinct conversion statuses (*BLO* 16 and *BLO* 22), because the exact position of trees within the plot had not been recorded in the validation dataset, we attributed the conversion status of the stand with the highest overlap to all trees of the plot. For each set of predicted probabilities, we computed two metrics: the scatter index, defined as the root mean squared error (RMSE) of prediction between predicted probability and observed trunk-base rot-hole presence-absence across all the trees of the dataset divided by the mean trunk-base rot-hole presence-absence, and the deviance of the model on the validation dataset. We thus obtained 3200 values of scatter index and deviance for the two models. As a benchmark, we also computed the values of scatter index and deviance for a null model where all the trees in the 2022 dataset had the same probability of harbouring a trunk-base rot-hole, set equal to the observed prevalence of trunk-base rot-hole in the 2022 dataset.

## 3 Results

### 3.1 Sampling effort

Operator pairs were able to survey 2 plots per day of fieldwork on average. The overall study thus took c.a. 17 operator.days. In 2021, 1462 oak trees were described over the nine surveyed plots. In 2022, 971 oak trees were described over the seven surveyed plots. The data has been made available on a public repository [Laroche, 2024]. In the 2021 calibration dataset, oaks of plots *BLO* 12 and *BLO* 17 harboured markedly lower DBH values than those of other plots (Figure 3A). This was expected since, in these plots, the final cut occurred less than 60 years ago (Figure 2), indicating that they are at an early stage of a regular high forest silvicultural cycle. Globally there seemed to be a positive relationship between oak trees DBH and their propensity to harbour a trunk-base rot-hole (Figure 3B).

**Figure 3:**
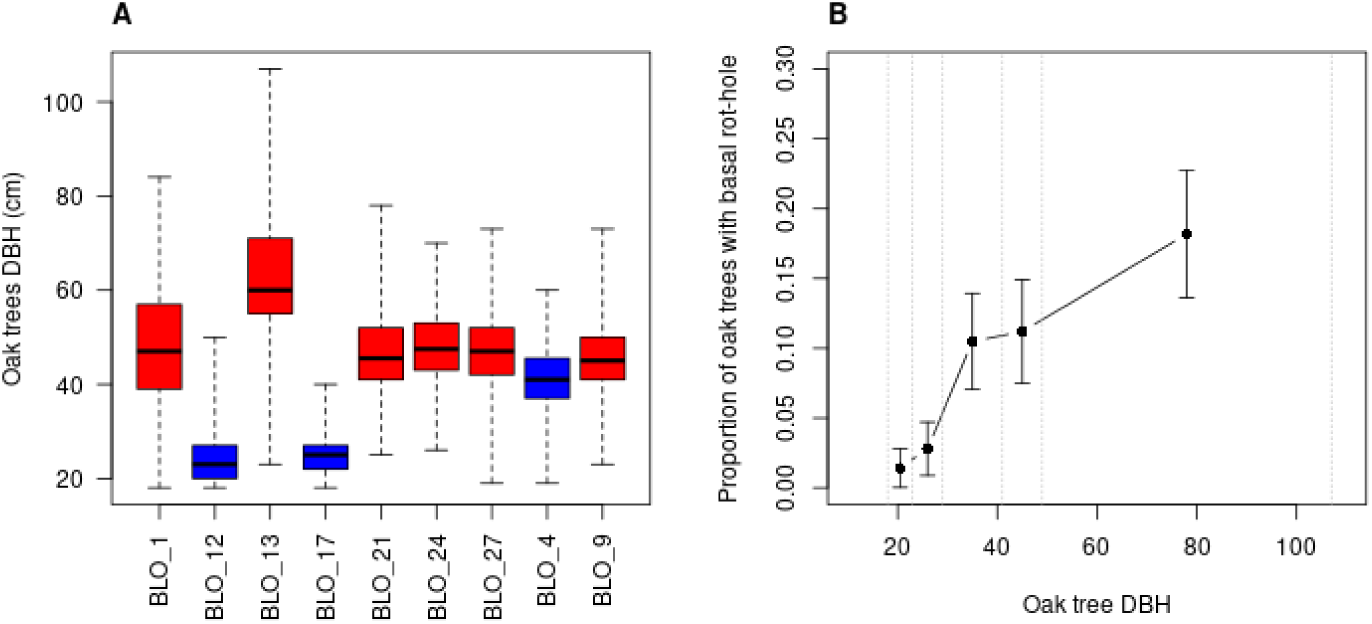
Summary of oak trees DBH and trunk-base rot-hole presence observed during the 2021 survey Grésigne. Panel A: distribution of oak trees DBH within each plot, boxes show extreme values, quartiles and median. Red boxes correspond to plots undergoing conversion, blue boxes correspond to plots where conversion was complete. Panel B: proportion of oak trees bearing a trunk-base rot-hole in five DBH categories. Categories were delineated using quantiles 0.2, 0.4, 0.6 and 0.8 of the DBH distribution. Borders of the bins are shown with vertical dotted grey lines. The proportion of trees bearing a trunk-base rot-hole is indicated in each bin using a dot. Bars indicate an asymptotic 95% confidence interval of the proportion.

The MCMC algorithm used to sample the posterior distributions of parameters yielded 3200 samples in 109 minutes for the non-informative priors and 3200 samples in 51 min for the informative priors using a conventional laptop (eight 1.60GHz processor units). An estimation summary is provided in Table 1.

**Table 1:**
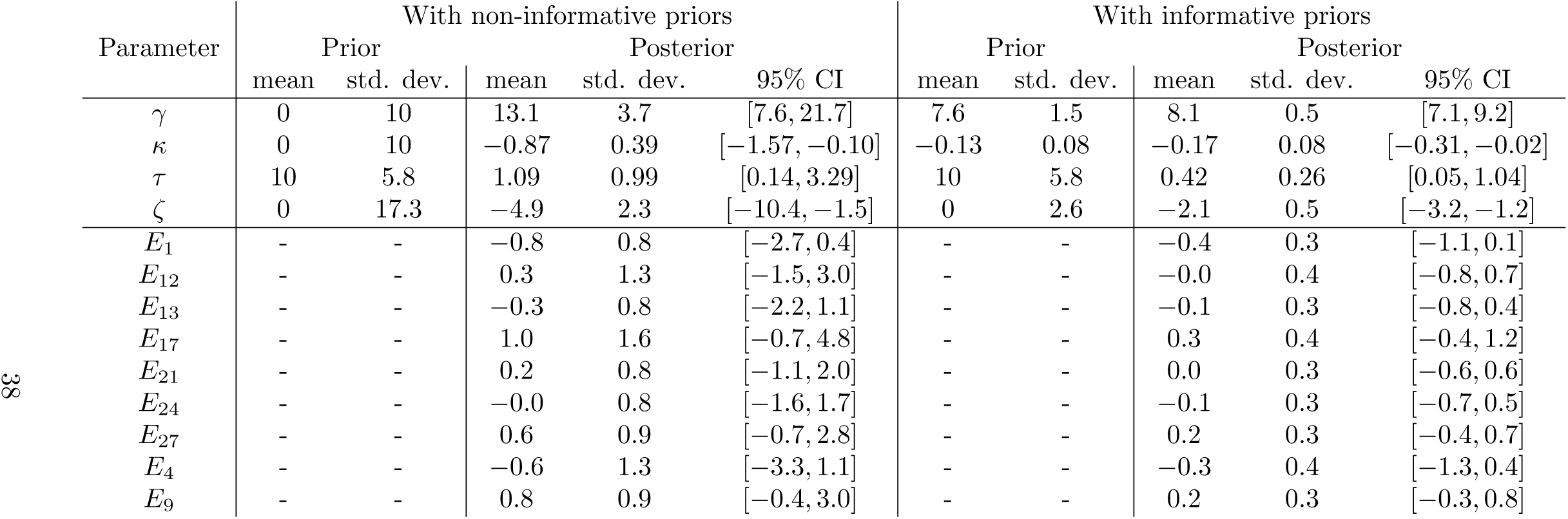
Summary of estimation of model of trunk-base rot-hole occurrence. As a reminder, *γ* is the log scale parameter of trunk-base rot-hole occurrence model in stands where conversion has ended, *κ* is the log shape parameter of trunk-base rot-hole occurrence model in all stands, *τ* is the standard deviation of plot random effect on log scale trunk-base rot-hole occurrence model and *ζ* is the effect on the log scale of trunk-base rot-hole occurrence model when moving from a stand where conversion has ended to a stand where conversion is ongoing. Pearson correlation between *γ* an *κ* informative prior distributions equalled 0.18. Pearson correlation between *γ* and *ζ* prior distributions equalled −1.5/√3 ≈ −0.87 informative cases. 3 *≈ −*0.87 for both non-informative and informative cases.

### 3.2 Model fit with non-informative priors

#### Posterior predictive checks

— The occurrence model of trunk-base rot-holes calibrated on the 2021 dataset with non-informative priors yielded a correct fit of the number of cavities in each plot, the total number of cavities on large oaks (DBH above 35 cm) and the total number of cavities on small oaks (DBH below 35 cm; Fig. S1).

#### Scale estimates in both conversion statuses

— The posterior distributions of parameters *γ* and *ζ* were highly variable (Tab. 1). The joint posterior distribution of *γ* and *θ* visually followed a curve suggesting low identifiability across a wide range of parameters (Fig. 4). The same phenomenon occurred between *γ* + *ζ* and *θ*. Those areas of the parameter space with low identifiability matched well the lines of non-identifiability predicted from equation (4). The posterior 95% credibility interval of the conversion effect *ζ* was broad but comprised only strictly negative values (Tab. 1). Comparing trees with the same DBH, occurrence probability of trunk-base rot-holes thus tended to be significantly higher in stands where conversion is still ongoing compared to stands where conversion has ended (Fig. 5A). As an illustration, the model predicted that a tree with DBH equal to 80 cm had 21% chance of harbouring a trunk-base rot-hole in plots undergoing conversion but only 3% chance in plots where conversion has ended.

**Figure 4:**
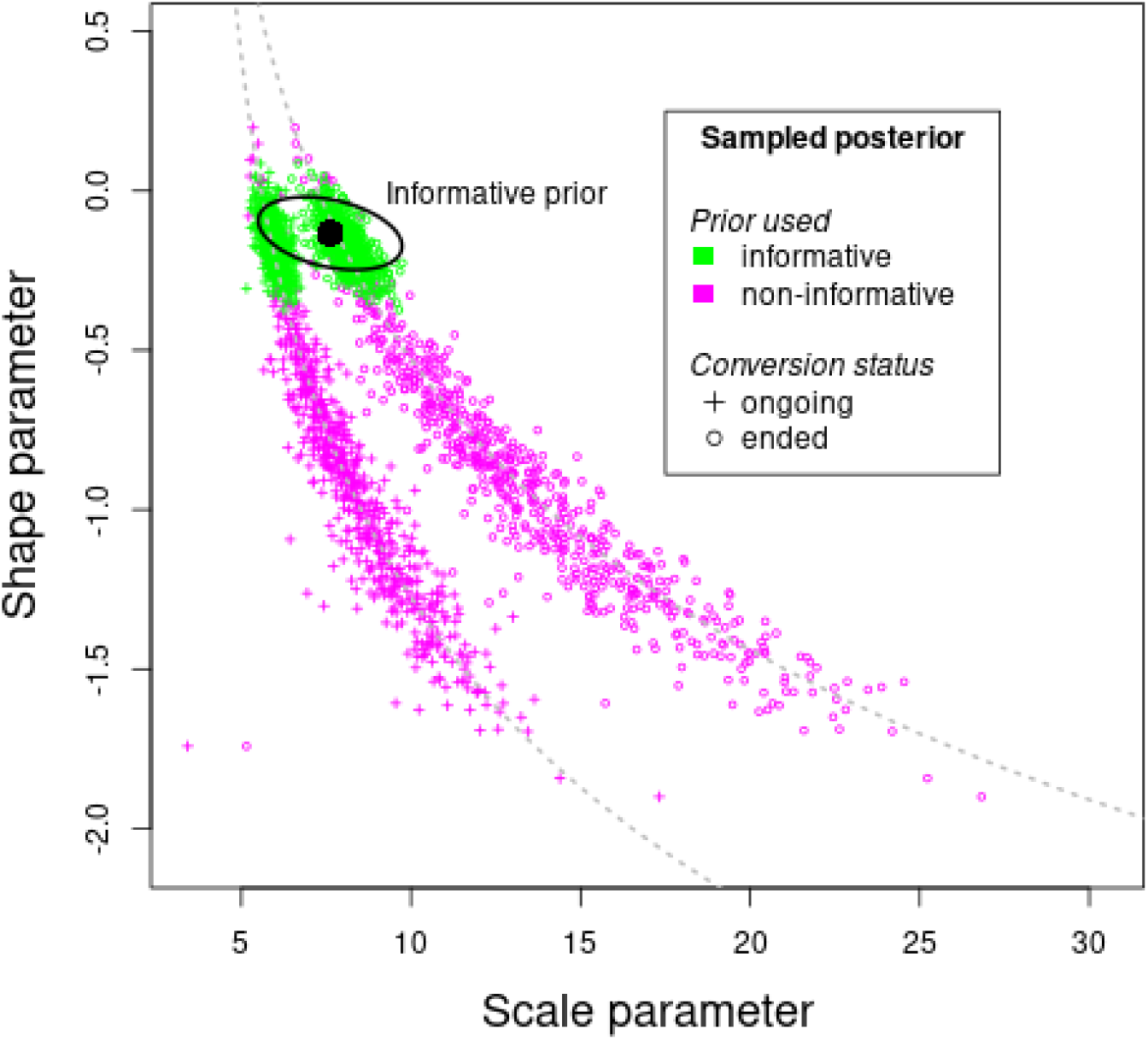
Sampled posterior distributions of shape and (log) scale parameters of the local model of trunk-base rot-hole occurrence of Grésigne calibrated on 2021 survey. For each sampled set of parameters, we reported a dot at coordinates (*γ, κ*), showing shape and scale parameters in stands where conversion has ended, and a cross at coordinates (*γ* + *ζ, κ*) showing shape and scale parameters in stands where conversion is ongoing. Sampled posteriors are derived either with non-informative (pink) or with informative (green) priors. A random subset of 20% of posterior samples is shown to improve readibility. The informative priors distribution is presented in black (black dot: mean of informative prior distribution; ellipse: one standard deviation to the mean). This distribution is the same in plots where conversion has ended (log scale parameter *γ*) and in plots where conversion is ongoing (log scale parameter *γ* + *ζ*). The grey dashed lines correspond, from left to right, to lines of non-identifiability for (*γ* + *ζ, κ*) and (*γ, κ*) respectively, as depicted in equation (4) of main text.

**Figure 5:**
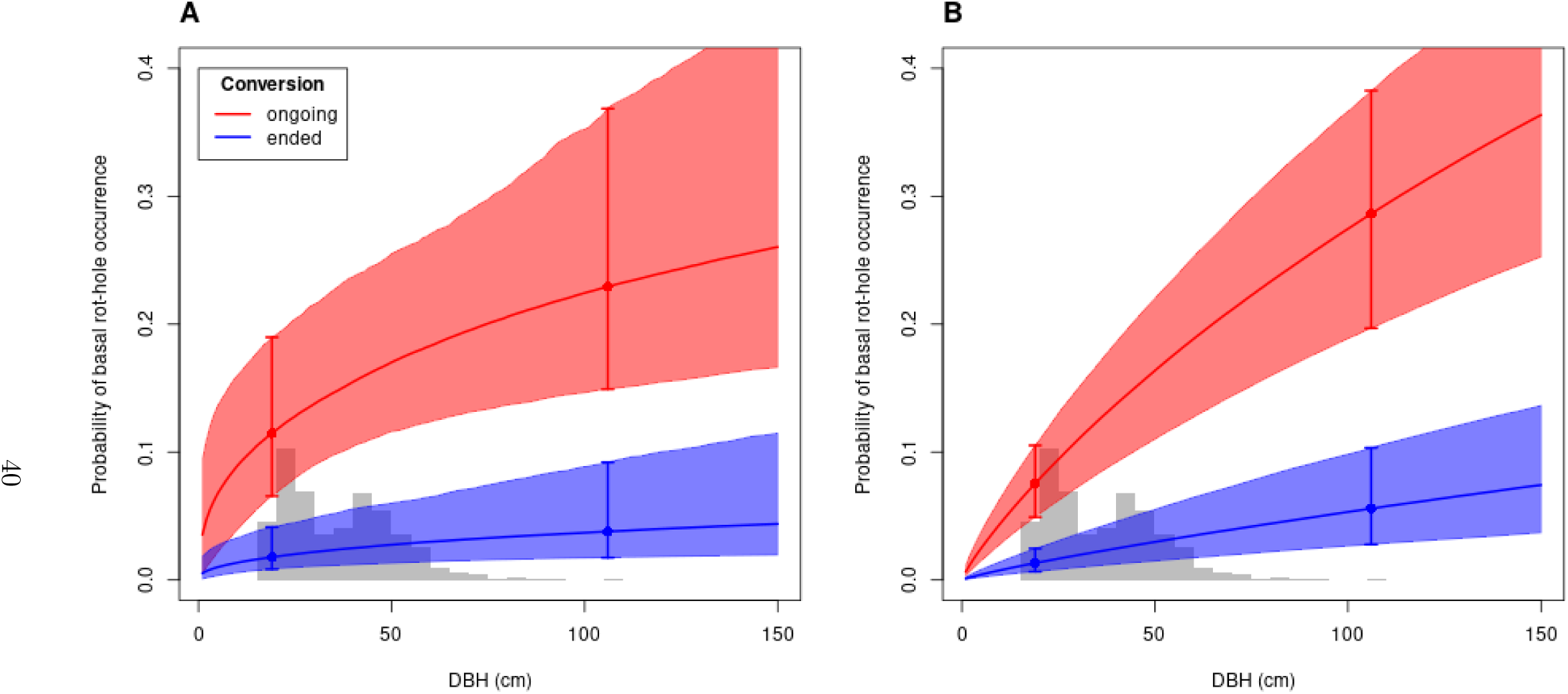
Probability of occurrence of a trunk-base rot-hole on Grésigne oaks as tree diameter increases, calibrated on the 2021 survey with non-informative (panel A) or informative (panel B) priors on parameters. Red color codes for plots undergoing conversion, blue color codes for plots that have ended conversion. The thick line indicates the mean probability of occurrence across the sampled posterior distribution of parameters, the envelope indicates the 95% confidence interval of the probability, empirically estimated from the sampled posterior distribution. Dots and error bars indicate the mean probability and associated confidence interval for the largest and smallest tree diameter values in the 2021 dataset. The grey histogram in the background recalls the distribution of oak DBH in the dataset.

#### Variation of trunk-base rot hole probability of occurrence with DBH

— Although the structure of the local model of trunk-base rot-holes occurrence necessarily implies that the probability of trunk-base rot-hole occurrence increases with tree DBH, we found that, for both conversion statuses, the probability harboured a quickly saturating profile. As a result, the probability of trunk-base rot-hole occurrence did not increase significantly across the range of DBH observed in our survey (see error bars and envelopes breadth in Fig. 5A). The 95% credibility interval of *κ* parameter contained only negative values, suggesting that the rate of trunk-base rot-hole creation on non-bearing oak trees decreased with tree diameter: large non-bearing trees would tend to develop trunk-base rot-holes more slowly per diameter growth unit than smaller trees.

### 3.3 Model fit with informative priors

#### Posterior predictive checks

— The occurrence model of trunk-base rot-holes calibrated with the informative priors on the 2021 survey yielded a correct fit of the number of cavities in each plot, the total number of cavities on large oaks (DBH above 35 cm) and the total number of cavities on small oaks (DBH below 35 cm; Fig. S2).

#### Scale estimates in both conversion statuses

— The posterior distribution of *γ* (log scale parameter in stands where conversion has ended) was located around the same values as the corresponding informed prior distribution (Fig. 4; Tab. 1), but was more narrow. The 95% credibility interval of the conversion effect *ζ* only included negative values. The posterior distribution of *γ* + *ζ* (log scale parameter in stands where conversion is still ongoing) departed from the informative prior distribution, being markedly shifted towards lower values (Fig. 4). Trunk-base rot-holes thus tended to occur at lower tree DBH in stands where conversion is still ongoing, compared to plots where conversion has ended (Fig. 5B). As an illustration, the model predicted that a tree with DBH equal to 80 cm had 23% chance of harbouring a trunk-base rot-hole in plots undergoing conversion but only 4% chance in plots where conversion has ended.

#### Variation of trunk-base rot hole probability of occurrence with DBH

— For both conversion statuses, the probability of occurrence of a trunk-base rot-hole harboured a concave but steadily increasing profile as DBH increased, irrespective of conversion status (Fig. 5B). The increase was faster in stands where conversion is still ongoing, resulting in a significant increase in trunk-base rot-hole probability of occurrence across the range of tree DBH observed in our survey. The posterior distribution of *κ* slightly shifted towards negative values compared to the informative prior distribution (Tab. 1). The posterior 95% credibility interval of *κ* parameter contained only negative values which suggested that the rate of trunk-base rot-hole formation on non-bearing oak trees decreased with DBH in Grésigne.

### 3.4 Comparing models

Using the informative priors improved the precision of the posterior sampled distribution of parameters compared to the model with non-informative priors (Tab. 1; Fig. 4). Quantitatively, the standard deviation of *κ* (resp. *γ*) posterior distribution was reduced by a factor of c.a. 5 (resp. 7; Tab. 1) between models with non-informative and informative prior.

Using informative priors instead of non-informative priors lead to lower scale parameters posterior distributions, and to a posterior *θ* distribution closer to 0. From the informative priors perspective, trunk-base rot-holes were therefore expected to occur at lower tree diameter and the pattern of decreasing rate of trunk-base rot-hole formation with tree growth seemed weaker.

The DIC was 749.6 for the model with non-informative priors and 750.9 for the model with informed priors. Therefore, the use of informed priors did not induce a significant increase in DIC compared to non-informative priors, suggesting no degradation of the fit when moving to informative priors. Both models calibrated with non-informative and informative priors seemed to show a lower prediction error on the 2022 validation dataset than the null model of trunk-base rot-holes occurrences among trees (Figure 6). However, the difference was significant at a 0.05 level only for the model with informative priors, irrespective of the metric considered. In addition, the model calibrated with informative priors showed a significantly lower mean prediction error than the model calibrated with non-informative priors, both in terms of RMSE and deviance.

**Figure 6:**
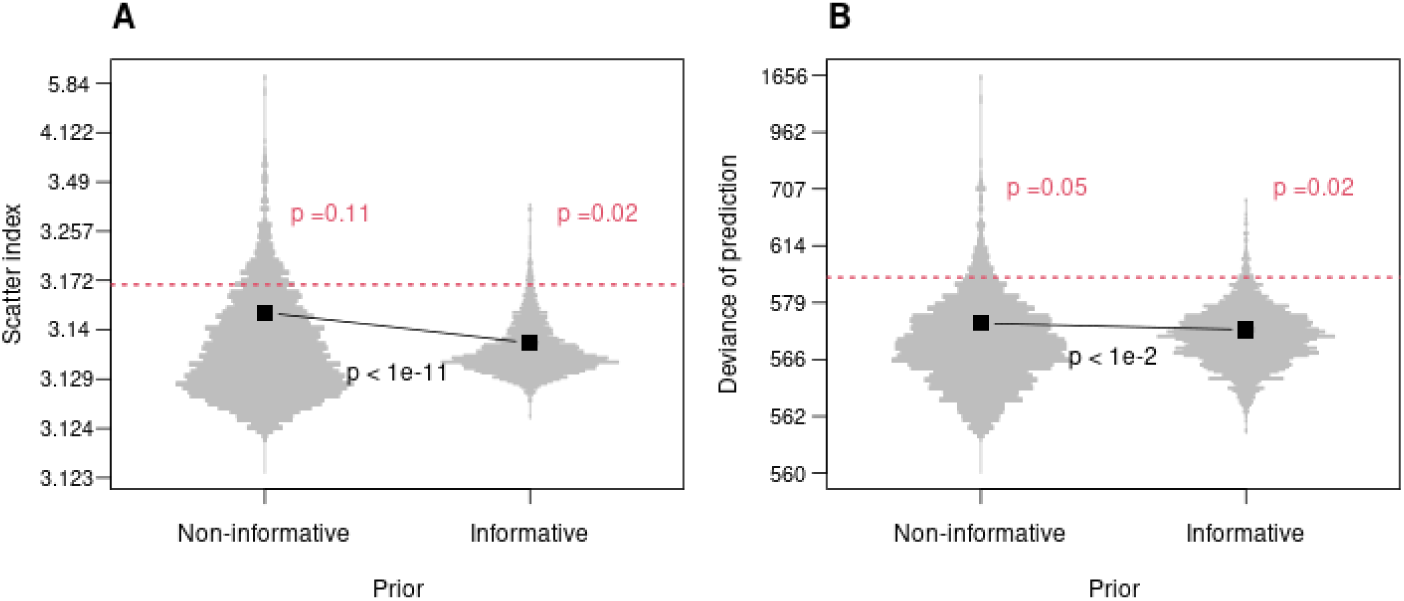
Prediction error of models with non-informative or informative priors, assessed on trunk-base rot-hole presence in the 2022 tree dataset. Prediction error is quantified as the scatter index (panel A) or the deviance of models (panel B). Mind the log-scale with offset on the y-axis. In both panels, for each prior type, the distribution of prediction error values associated to the sampled posterior distribution of parameters is presented using a violin plot. The mean prediction error is pictured with a black square. Mean prediction errors are compared using an asymptotic test, and the resulting p-value is reported in black. The red horizontal dashed line presents the predictive performance of the null model of trunk-base rot-holes among trees in the 2022 tree dataset. The prediction errors of models with non-informative or informative priors are compared to this benchmark using a one-sided test and the resulting p-value is reported in red.

## 4 Discussion

We explored whether the use of informative priors could improve the identifiability, the precision of estimates and the predictive abilities of a model of trunk-base rot-holes occurrence in Grésigne forest — a context where stand management and tree DBH were confounded. Without prior information, the low variance of tree DBH within management modalities rendered the model poorly identifiable and prevented the detection of an effect of tree DBH *per se* across the range of tree DBH explored. By contrast, using informative priors contributed to improve the precision of estimates and lead to detecting a positive effect of tree DBH *per se* within stands where conversion is ongoing. Informative priors did not degrade the model fit and clearly improved predictive abilities on a novel dataset. In particular, while the model without prior information did not lead to a significantly better prediction of trunk-base rot-holes occurrence than a purely random guess, the model with informative priors did. Irrespective of the prior used, models suggested that the high recruitment of trunk-base rot-holes in Grésigne may be a temporary management effect in stands undergoing conversion from coppice-with-standards to high forest through sprout thinning, which will lead to conservation issues for cavicolous saproxylic species when all conversions are complete. Below, we further discussed each of these points in turn.

### Confounded stand management and tree DBH lead to low identifiability and no DBH effect *per se* without prior information

— Because stand management modalities and tree DBH are confounded in Grésigne forest, the variance of tree DBH was low within management modalities, which generated several issues in the model with non-informative priors. First, it lead to weak identifiability : an infinite subset of parameter values for the shape and scale parameters of trunk-base rot-holes occurrence model yielded similar likelihood values, hence forming the lines of low identifiability observed in results. As we saw, this induced low precision of estimates. It also raised a more profound issue of posterior sensitivity to the non-informative prior choice. Because of these lines of low identifiability, the posterior distribution does not converge as the variance of the non-informative prior increases, and the wider the vague prior used on parameters, the larger the positive bias on posterior estimates. These issues have been extensively discussed by Gelman [2006], and our study followed the advice of this author of setting moderately large variance in our non-informative priors. Second, the confounded stand management and tree DBH effects hindered the detection of an effect of tree DBH *per se* on trunk-base rot-holes occurrence, which seemed only affected by management modalities. Finding an effect of management was expected, because of the particular history of the Grésigne forest and more generally because management practices are known to be a strong driver of tree-related microhabitats occurrence [Vuidot et al., 2011]. By contrast, the absence of tree DBH effect was surprising, and most likely due to a lack of statistical power. A trunk-base rot-hole can show various development stages, but it is likely to last as long as the bearing tree persists [Larrieu et al., 2022], contrary to other TreMs that can disappear or evolve to new types [Spînu et al., 2024]. Therefore, the probability of observing a trunk-base rot-hole on a tree must increase in time. Because the DBH of a tree also increases in time, one expects a positive correlation between DBH and trunk-base rot-hole occurrence, as observed in previous empirical studies [Courbaud et al., 2017, 2022, Kozák et al., 2023]. The saturating profiles of the occurrence probability predicted by the model with non-informative priors rather suggested that most trunk-base rot-holes do not develop slowly with tree growth, but rather appear on a random fraction of trees before they reach the 17.5 cm DBH threshold at which our observations started. It should be noted that this scenario might not be completely unrealistic in Grésigne stands where conversion towards high forest is still ongoing because, in high forest generated from sprout thinning, the tree stump (where the trunk-base rot-holes occur) can be older and larger than the stem which is actually measured. This could explain the apparent decoupling between tree DBH and trunk-base rot-hole occurrence, and why the model with non-informative priors predicted that the rate of appearance of trunk-base rot-holes decreases with tree growth. Gouix et al. [2015] illustrated this effect in Grésigne forest by showing that the diameter at 30 cm height from the ground explained better the presence of the violet click beetle (*L. violaceus*) in trunk-base rot holes than the DBH of stems. However, this sprout-thinning effect cannot explain the absence of relationship between DBH and trunk-base rot-hole occurrence in stands where conversion has ended, which are managed as regular high forest.

### Informative priors overcame the issues raised by confounded tree DBH and stand management

— Introducing informative priors overcame the identifiability issue mentioned earlier by forcing shape and scale parameters of the trunk-base rot-hole occurrence rate function to lie in a compact region of the parameter space resulting from a compromise between the informative priors influence and the lines of low identifiability. This forcing restored the positive relationship between the probability of trunk-base rot-hole occurrence and tree DBH which was expected from a temporal process of appearance. However, given their strong impact on the analyses, one must carefully consider whether the informative priors used were relevant to the particular context of Grésigne forest. Having adopted a transparent, reproducible methodology to obtain these priors was an essential pre-requisite to discuss this point [Dennis, 1996, McCarthy and Masters, 2005]. Informative priors were obtained from a continental study where the management modalities considered here (ongoing conversion or conversion ended) were not distinguished: they both fell in the broad ‘managed’ category, along with other sylvicultural practices [Courbaud et al., 2022]. The informative prior on the scale parameter derived from the continental model may thus be more or less adapted to the local management modalities in Grésigne, depending on whether these modalities are common or rare in the continental dataset, potentially leading to biased estimates. The precise management modalities of forests used in the continental dataset were not available, but one could conjecture that the conversion through sprout thinning that is ongoing in a part of Grésigne is quite specific compared to classic management practices, and that the continental network of forest sites used to build the informative priors would therefore be little representative of those stands where conversion is ongoing, but better adapted to stands where conversion has ended.

### Accounting for local management specificities when using informative priors was a key step to limit estimation biases

— Contrary to what could be expected from the atypical conversion history of Grésigne, the use of informative priors did not induce a significant decrease in the fit, suggesting no estimation bias. The absence of bias was further corroborated by the significant increase in the ability to predict trunk-base rot-holes occurrence in novel forest stands of Grésigne. The estimation bias remained limited because our local model allowed for a difference in trunk-base rot-holes occurrence rate between stands undergoing conversion and stands where conversion has ended. Thanks to this decoupling, the posterior distribution of the scale parameter driving trunk-base rot-holes occurrence rate in stands under conversion could depart from the informative prior, while the scale parameter in stands managed as classic high forest could remain close to the informative prior. Stands undergoing conversion in Grésigne harboured atypically high trunk-base rot-holes occurrence compared to other contexts encountered at continental scale, as stressed by the monography of Ducasse and Brustel [2008]. By contrast, stands where conversion has ended harboured more classic rot-holes occurrence rates, within the range of what can be observed at continental scale. This marked difference between both management modalities clearly suggested that if the heterogeneity of stand management in Grésigne had been overlooked in the model design, informative priors would likely have lead to strong biases in the estimation of trunk-base rot-holes accumulation process. This warns against using informative priors on TreMs dynamics in a standardized and systematic way without careful case-by-case adaptations taking into account the specificities of the local context with respect to previous studies that were used to build the prior. Otherwise, one expects the same procedure to generate unpredictable biases across case studies (as illustrated in another forest topic by Morris et al. [2015]).

### No tractable alternative tree sampling strategy could have decoupled stand management and tree DBH better in Grésigne

— Another way to overcome the issues raised by the correlation between tree DBH and stand management could have been to actively design a sampling strategy that would decouple both covariates, i.e. that would aim at generating similar DBH distributions within the two management modalities studied here. This was not possible with a plot-based sampling strategy in Grésigne. Plots tend to extract representative distribution of DBH from underpinning stands, and stand management and tree DBH are confounded. Therefore, in Grésigne, we expected that any plot-based strategy would generate confounded stand management and tree DBH covariates. Overcoming the confounding effect through sampling would therefore have implied to drop, at least partially, the plot-based sampling strategy and e.g. sample individual trees across the forest. This would have allowed for actively sampling large oak trees in stands where conversion has ended and small oak trees in stands where conversion is ongoing, hence increasing the tree DBH variance in both management modalities and homogenizing their distribution. However this would have raised several issues: (i) the standing variance of tree DBH within stand management modalities may not have been sufficient to implement this strategy, (ii) the ‘outlier’ trees included in the sample may have been very atypical and little representative of the trunk-base rot-hole accumulation process; (iii) such an individual-based sampling strategy would have considerably increased the time spent per sampled tree. Using informative priors thus seemed a more tractable option.

### The high recruitment of trunk-base rot-holes in Grésigne is a tempo-rary management effect

— Trunk-base rot-holes were more frequent and occurred earlier with tree growth in stands undergoing conversion from coppice-with-standards to high forest compared to stands managed as classic high-forest. This effect was sufficiently strong to be detected even with non-informative priors. While sprout thinning is the major driver of trunk-base rot-hole formation in the former situation (Fig 1A), only weaker factors remain active in classic high forest, mostly damages related to logging and skidding (Fig 1B). These findings raised a conservation issue. Since classic high forest treatment is not generating trunk-base rot-holes at the same rate as the conversion regime, the trunk-base rot-holes renewal will necessarily decrease as more stands end their conversion, and the Grésigne forest may not remain a hotspot of trunk-base rot-holes in the long term. Also, trunk-base rot-holes generated by logging damages on trees may be of smaller size (a critical feature for saproxylic beetles; Gouix et al. [2015]) and lower quality than holes induced by sprout thinning in terms of habitat for rot-hole dwelling species. We did not explore this qualitative dimension in our study, although more elaborated models of trunk-base rot-holes development stages [Larrieu et al., 2018] with tree growth could be developed in this direction. A reasonable integrative management action in line with our results would thus be to keep oak trees starting back from natural sprouting disseminated within the new stands, to ensure the recruitment and the connectivity of future trunk-base rot-holes, as suggested by Gouix [2011].

### Using informative priors was simple and beneficial in our study, and should thus be further explored in local applied contexts

— With 17 operator.days of fieldwork (the effort needed to collect the 2021 calibration dataset) and less than one hour of computing on a conventional laptop, we have been able to derive a model of trunk-base rot-hole occurrences combining local data and informative priors which yielded better estimation and prediction than a model based on local data only. The increase of prediction abilities was admittedly moderate in terms of magnitude, as could be expected from a rare tree-related microhabitat like trunk-base rot-holes. However, contrary to the model with non-informative priors, the informed model was able to outperform a purely random model of trunk-base rot-hole occurrence distribution among trees, which is an important qualitative achievement to establish the interest of the modelling approach. Such model could now be coupled with management scenarios (e.g. in forest simulation platforms; Dufour-Kowalski et al. [2012], Courbaud et al. [2015]) to design viability pathways for trunk-base rot-holes and associated biodiversity as the renewal of the forest goes on. Our model will nonetheless need re-evaluation in the future, for climate-driven changes in growth regime and mortality may affect the relationship between diameter and trunk-base rot-hole presence, especially in Grésigne forest where sessile oaks are at the edge of their environmental niche and face low-fertility conditions that make them particularly vulnerable to climate changes.

## Acknowledgements

FL thanks Y. Grzelec, the head of Grésigne forest management unit for his help in planning the fieldwork, A. Ardanuy-Gabarra, A. Brin, W. Heintz, S. Ladet, J. Molina and G. Parmain for their help during the field work. All the authors thank M. Legay, J. Archambeau, G. Sommeria-Klein and S. Schmitt for their insightful comments and feedbacks during the peer-reviewing process. A CC-BY public copyright license has been applied by the authors to the present document and will be applied to all subsequent versions up to the Author Accepted Manuscript arising from this submission, in accordance with the grant’s open access conditions.

## Data, scripts, code, and supplementary information availability

Data used in the study has been made available on a public online repository [Laroche, 2024]. Statistical scripts have been made available on a public online repository [Cottais and Laroche, 2024]. Supporting information has been made available on a public online repository [Laroche, 2025].

---

**Fig. S1:**
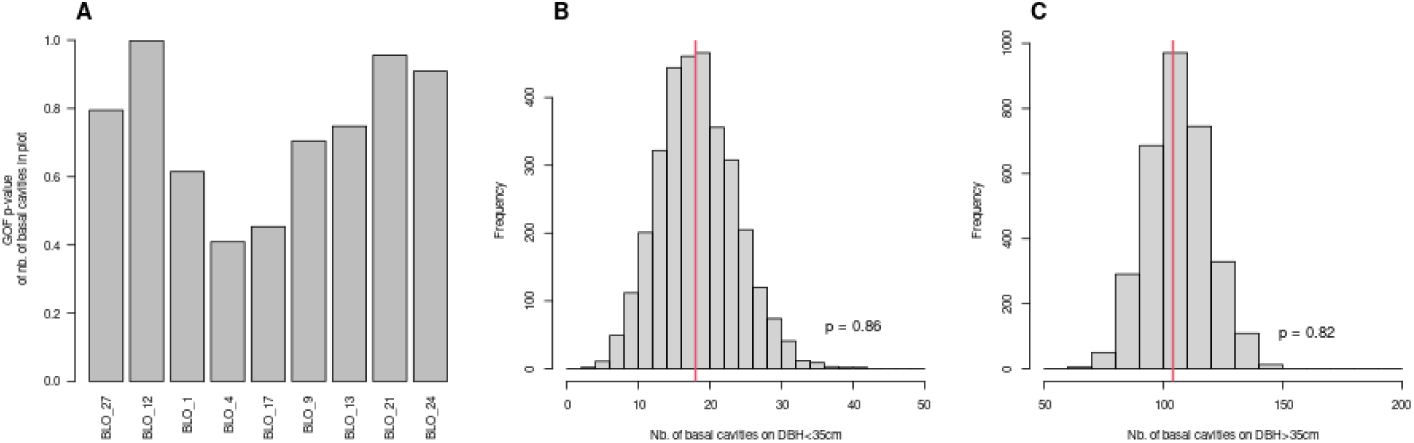
Goodness-of-fit (GOF) of the occurrence model for trunk-base rot-holes of Grésigne with non-informative priors, calibrated on 2021 survey. Panel A: GOF p-value of a bilateral test for the number of trunk-base rot-holes in each plot. Panel B and C: simulated (grey histogram) versus observed (red bar) number of trunk-base rot-holes on trees with DBH below 35 cm (panel B) and above 35 cm (panel C) over the 2021 survey. In panel B and C, the GOF p-value associated to the red bar is reported directly on the graph.

**Fig. S2:**
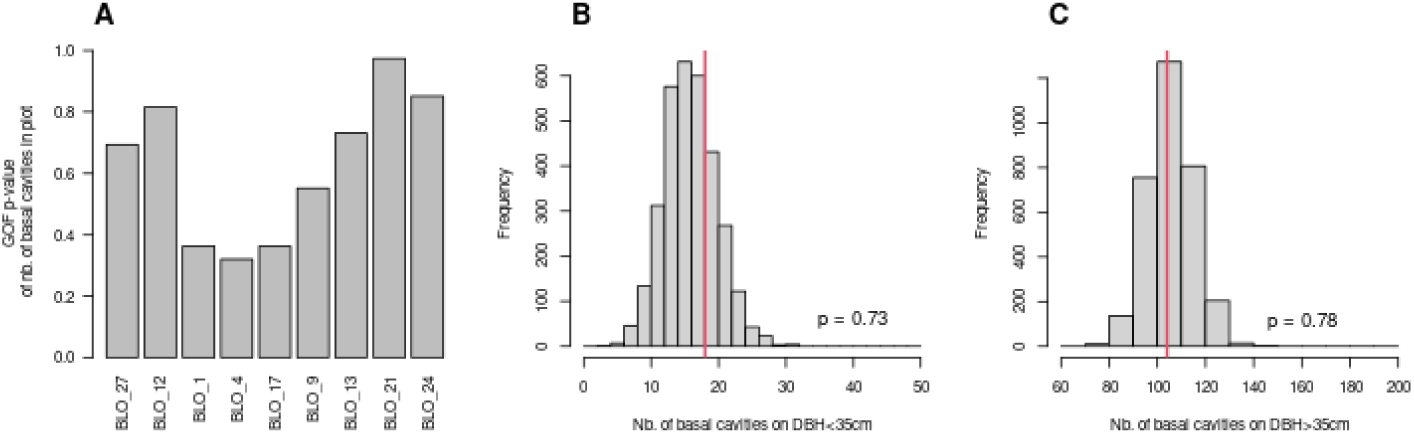
Goodness-of-fit (GOF) of the occurrence model for trunk-base rot-holes of Grésigne with informed priors calibrated on 2021 survey. Panel A: GOF p-value of a bilateral test for the number of trunk-base rot-holes in each plot. Panel B and C: simulated (grey histogram) versus observed (red bar) number of trunk-base rot-holes on trees with DBH below 35 cm (panel B) and above 35 cm (panel C) over the 2021 survey. In panel B and C, the GOF p-value associated to the red bar is reported directly on the graph.

